# Threshold public goods game in well mixed populations: Cooperation in density independent and density dependent cases

**DOI:** 10.1101/2023.09.26.559593

**Authors:** Adél Károlyi, István Scheuring

## Abstract

The threshold public goods game is one of the best-known models of nonlinear public goods dilemmas. Cooperators and defectors typically coexist in this game when the population is assumed to follow the so-called structured deme model. In this paper we develop a dynamical model of a general N-player game in which there is no deme structure: individuals interact with randomly chosen neighbours and selection occurs between randomly chosen pairs of individuals. We show that in the deterministic limit the dynamics in this model leads to the same replicator dynamics as in the structured deme model, i.e. coexistence of cooperators and defectors is typical in threshold public goods game even when the population is completely well-mixed. We extend the model to study the effect of density dependence and density fluctuation on the dynamics. We show analytically and numerically that decreasing population density increases the equilibrium frequency of cooperators till the fixation of this strategy, but below a critical density cooperators abruptly disappear from the population. Our numerical investigations show that weak density fluctuations enhance cooperation, while strong fluctuations suppress it.

**Highlights:**

- The dynamics of threshold dilemma game in a well-mixed population.
- Fluctuation of density can help or suppress cooperation.
- Cooperation is more pronounced at lower density.
- Cooperation abruptly disappears below a critical density level.

## 1. troduction

When an individual performs an act that is costly to itself but beneficial to its conspecifics (e.g. guarding, feeding, mobbing predators), this is called altruistic behaviour (West et al. (2007)). Similarly, when costly cooperation between two or more individuals results in something that benefits them (e.g. group hunting, alloparental care), this is called cooperation (or mutual altruism). These altruistic and cooperative interactions are very common in biology. Their presence is crucial to the success of a species in a given habitat, and in many social species the whole population would be unsustainable without cooperation between individuals, just think of ants, termites, cooperatively breeding birds, and even Homo sapiens (West et al. (2021)). Moreover, the emergence of new forms of cooperation within newly emerged evolutionary units is an important feature of major evolutionary transitions (Maynard Smith and Szathmáry (1995)). There is therefore no doubt that such interactions have a very significant impact on the functioning of the biosphere as a whole. However, their evolutionary origin and stability is by no means a simple problem. Assuming that the altruistic or cooperative acts are costly, one can argue that these mutants cannot invade the population of non-cooperative individuals, making their evolutionary origin problematic. Furthermore, if we neglect the problem of origin and assume that each individual is already altruistic or cooperative, then the mutant cheater (defector) who doesn’t invest in the cooperative act will only benefit from the interaction at no cost, so it will spread in the population. Consequently, the evolutionary stability of this cooperative behaviour does not seem to be a trivial question.

Several solutions to this evolutionary contradiction have been proposed in recent decades. One of the most important mechanisms supporting the evolution of altruistic or cooperative behaviour is kin selection (Hamilton (1964)). The idea is that because of the kinship between the helper (altruist or cooperator) and the helped individual, the inclusive fitness of the altruist will also increase as a result of its act. In other cases, even when kinship is negligible, if altruists interact with each other preferentially compared to interactions with cheaters (positive assortment of altruists, mate choice) in the population, or if cheating is punished or cooperation is enforced, then the fitness of cheaters is depressed and cannot spread among helpers (West et al. (2021)). A similar, though in some aspects different, mechanism occurs when multi-level selection is at operation, i.e. individuals within a population temporarily form groups whose success depends on the quality (and/or quantity) of cooperation that occurs within them (Okasha (2006)). In such a case, even though helpers are at a disadvantage compared to cheaters, the counteracting effects of competition between groups and individuals lead to a stable equilibrium between helpers and cheaters (Wilson (1977); Okasha (2006)).

A biologically important subset of cooperation or mutual altruism is when (some) individuals create a public good from which all can benefit (West et al. (2021)). This is what happens when bacteria release degrading enzymes or toxins into the extracellular matrix (Patel et al. (2019a,b)), when in a herd of animals grazing in groups, some individuals watch for predators and signal to others when there is danger (Clutton-Brock et al. (1999)), or when predators hunt in a cooperative group (Bednarz (1988); MacNulty et al. (2011); Yip et al. (2008)). In these cases, a model that fits the phenomenon well is the nonlinear public goods game (NLPGG) where the public good distributed among the participants is a nonlinear function of the number of cooperators, which, according to experimental observations (e.g. Bednarz (1988); Clutton-Brock et al. (1999); Archetti and Pienta (1995); MacNulty et al. (2011); Rosenthal et al. (2018)) and theoretical considerations (e.g. Archetti and Scheuring (2016); Archetti and Pienta (2019)), typically follows a saturating sigmoid curve (Fig. 1 B). The general sigmoid curve can be approximated by a threshold function (Fig. 1 C), and it can be shown that the threshold public goods game (TPGG) behaves qualitatively in the same way as the sigmoid public goods game (SPGG) (Archetti and Scheuring (2012)).

**Figure 1.**
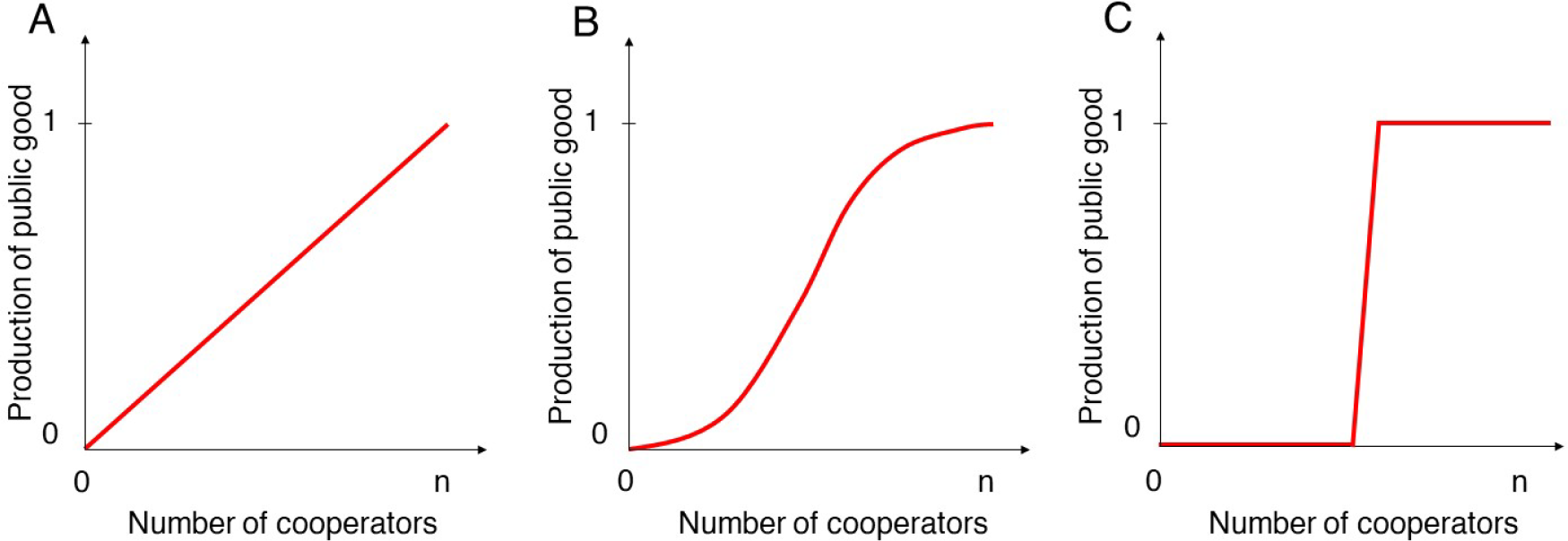
The level of public goods in function of number of cooperators in the interaction group. **A:** Traditional linear public goods game (PGG), **B:** Nonlinear public goods game with sigmoid benefit function (SPGG), **C:** Threshold public goods game (TPGG).

To study the SPGG or TPGG, the following model is used: There are two strategies, the cooperator (C) who invests in the public good and the defector (D) who does not. It is assumed that the population is very large and individuals are randomly assigned to local interacting groups of size *N* individuals where they share the public good. The replicator dynamics is determined by the average fitness of the cooperative and defective strategies which are computed as the weighted average of the fitnesses in the local interacting groups. It can be shown that there are four qualitatively different solutions of the dynamics, depending on the *k* and the maximal benefit and cost ratio (Archetti and Scheuring (2012) where the coexistence of cooperators and defectors is one of the common stable steady state of the dynamics (for more details see Fig 2. and in the section 3). This result clearly shows that cooperators and defectors can coexist stably, despite the absence of any spatial aggregation or partner selection, when the public good game is a threshold (or sigmoid) saturating function. However, the model and its interpretation can be criticised by arguing that there is a so-called structured deme population structure in the background (Fig. 3). That is, while there is no direct aggregation of cooperators while the interacting groups are formed, the success of cooperators and defectors, which determines the dynamics, is obtained as the weighted average of the success rates in these groups. So strategies are practically places into a multilevel selection situation (Matessi and Jayakar (1976); Wilson (1977); Charlesworth (1979); Okasha (2006)). Consequently, the stable coexistence of cooperators and defectors is not so surprising, since individuals in groups with enough cooperators send more offspring to the next generation than individuals with few or no cooperators. In addition, this also raises questions about the biological feasibility of the model, since multilevel selection is not thought to be widespread in Nature. Another weakness of the model, like many evolutionary game theoretical models, is that interactions are assumed to be density independent. To understand why this is a problem let us consider for example a population of bacteria that release an extracellular degrading enzyme. The efficiency of the enzyme, i.e. the benefit to the public good, depends not only on the frequency of the enzyme releasers (i.e. cooperators), but also on the density of cooperators in the habitat. There are, of course, some papers that examine the public good dilemma in the density-dependent case. In a keystone paper Hauert et al. (2006) (Hauert et al. (2006a)) studied the classical linear public goods game (PGG) (Fig. 1 A) in the density-dependent case. They worked in the framework of structured deme model where it is easy to show that in the density-independent case, the defector wins if the reward factor of a cooperator’s contribution (*r*) divided by the number of individuals (*N*) in the interaction group is less than one. If the opposite is true (*r*/*N* > 1), then the cooperators are the winners of the selection (the derivation is shown in section 2.5). They modified the model so that the lower fitness associated with the spread of defectors also reduces the density of the population, resulting in a smaller interaction group size *N*. Thus, if initially *r*/*N* < 1 (*r*/*N* > 1), then due to the spread of defectors (cooperators) density decreases (increases), and the population moves to the *r*/*N* > 1 (*r*/*N* < 1) state where the cooperators (defectors) begin to spread and thus the density increases (decreases). These opposite processes result the existence of an equilibrium point where cooperators and defectors coexist and *r*/*N* = 1. Depending on the model parameters, this fixed point can be stable or unstable. The dynamics is also determined by the initial frequency of cooperators, since if there are not enough cooperators in the population, it can die out. Together, these factors can lead to stable coexistence of cooperators and defectors (stable fixed point, limit cycle), fixation of cooperators or extinction of the whole population (Hauert et al. (2006a,b)). So, the behaviour of the density-dependent model differs significantly from that of the density-independent model. This highlights the importance of density dependence in N-person games.

**Figure 2.**
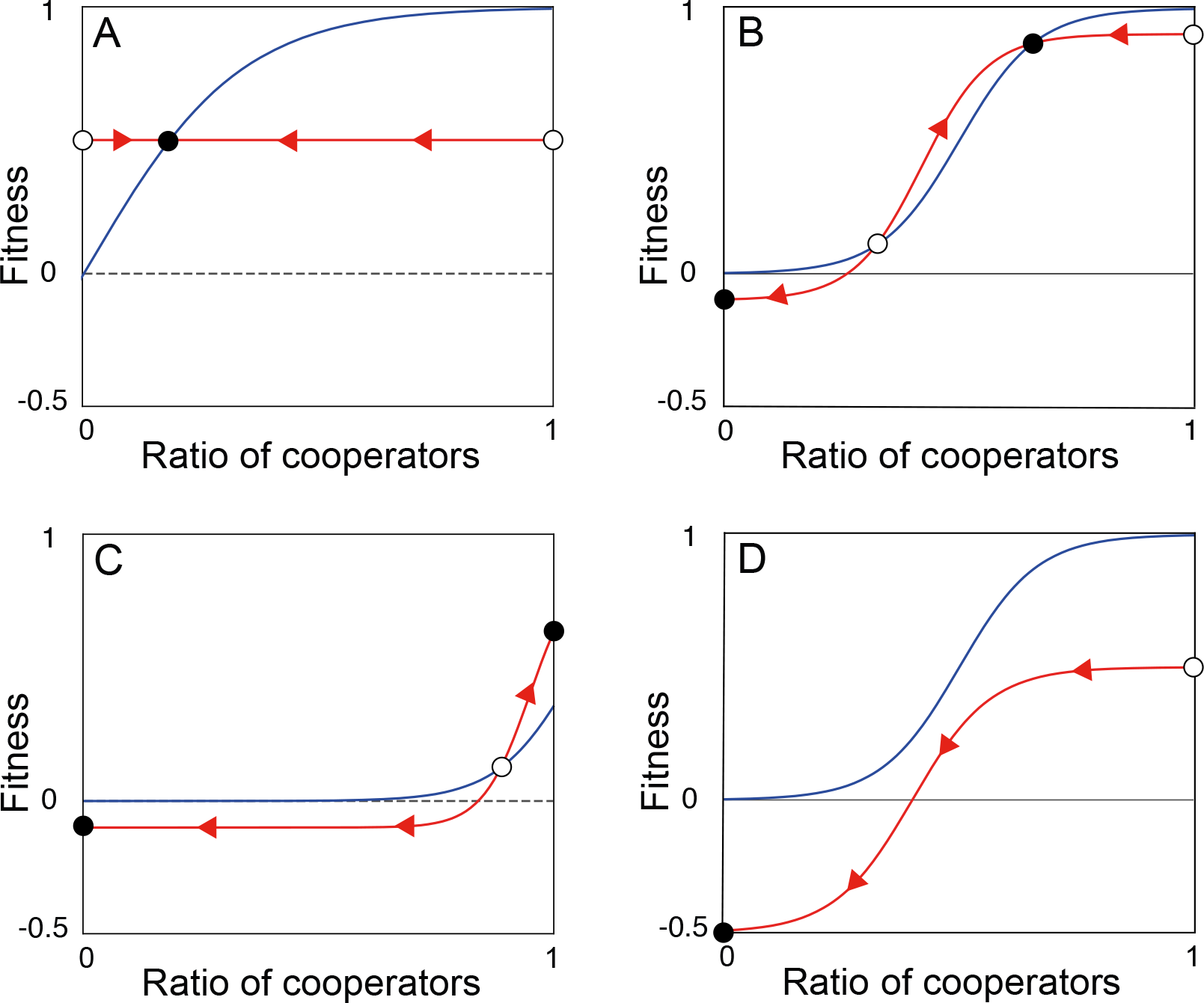
The graphical demonstration of the possible characteristically different dynamics of cooperators and defectors in the SPGG or TPGG by using structured deme model and replicator dynamics. The average payoffs (fitness) of defectors (blue line) and cooperators (red line) are depicted in function of the frequency of cooperators in the population. Empty circles denote the unstable fixed points while filled circles represent the stable fixed points of the dynamics. Arrows indicate how the frequency of cooperators change at different ratio of cooperators. bf A: *k* = 1 Cooperators and defectors stably coexist. **B**: 1 *< k < N*, the maximal marginal benefit of cooperation is above a critical level. There are two stable and two unstable fixed points. Cooperators coexist with defectors if the cooperators’ ratio is above a critical level initially. **C**: *k* = *N*, the maximal marginal benefit of cooperation is above a critical level. Depending on the initial ratio of cooperators either defectors or cooperators fixate in the population. **D**: 1 *< K ≤ N*, the maximal marginal benefit of cooperation is below a critical level. defectors win the selection.

**Figure 3.**
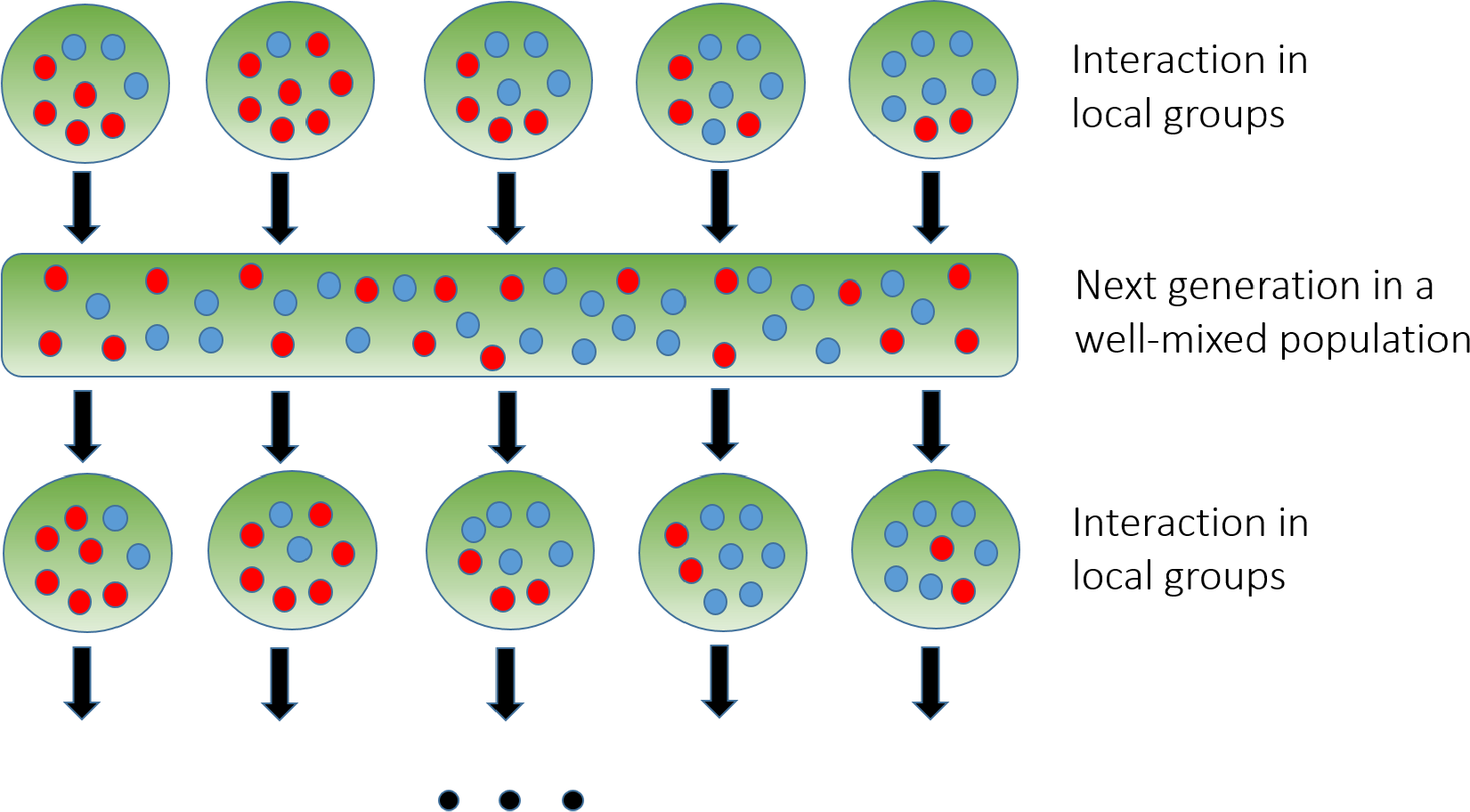
The schematic figure of the structured deme model. Interactions between individuals following different strategies (denoted by different colors) take place in local groups. The groups are randomly assembled from a large well-mixed population. The payoffs obtained in this phase determine the fitness of the individuals, which is used to send offspring to the next generation, where the population is again well-mixed. In the same way as in the previous step, local groups are then formed again randomly, taking into account the actual frequency of the strategies.

In this paper we aim to develop a dynamical model of a general N-player game in which there is no deme structure, only randomness in the composition of interacting strategies are taken into account. We show that the model leads to replicator dynamics identical to those of a population generated from the structured deme model. That is, in the case of TPGG, cooperators and defectors can coexist without multi-level selection. We also develop a density-dependent version of the TPGG model and analyze the dynamical behaviour of it. We study the stochastic agent-based version of the density independent and the density dependent models numerically, which also support our results from the deterministic analytical models. We also investigate the effect of fluctuations in the steady-state density in the framework of the stochastic agent-based model.

## 2. The general model

### 2.1. Basic assumptions

We consider a game *G* determined by the strategy set *S* and the payoff function *P*, i.e. a *G* = ⟨*S, P*⟩ game. The strategy set *S* contains a finite number of different pure strategies. Individuals follow one of these strategies and the game is symmetric. The population is very large, i.e. population size *K* ≫ 1, and individuals interact according to the game *G* in a local neighbourhood of size *N*. The population is well mixed, i.e. the *N* interacting individuals are randomly selected from the population without any selective sorting. We assume that each individual participates in the game exactly the same number of times in the so-called *interaction phase*. This phase is followed by the replication or *update phase*, where pairs of individuals are randomly selected from the population and the probability of replacing each other in the population is determined by their actual relative payoffs (Hilbe (2011)). It is important to emphasise that the competitive success of individuals in the replication phase is determined by the payoffs received in the game *G* by engaging with distinct local neighbours earlier. So there is no group formation and aggregation in the population, we just exploit the fact that interaction is local. The interaction and replication phases follow each other sequentially.

### 2.2. Notations

Let *n*_*i*_ be the number of individuals following strategy *i* in the given interaction group of size *N*, so 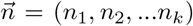 is the vector of these numbers, where 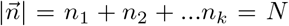 is the total number of individuals interacting with each other. The neighbourhood composition of a focal individual is denoted by 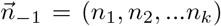, where 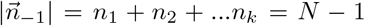, and the strategy composition in the whole group together with focal strategy *i* is 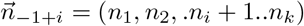.

An individual following strategy *i* receives a payoff 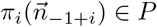 in the interacting phase. This notation emphasises that the payoff is determined by the neighbours and the actual strategy of the focal individual.

### 2.3. Imitation dynamics

After the interaction phase, we randomly select pairs of individuals from the population and compare their payoffs. Since the population is very large, the probability of selecting individual pairs from the same interacting group is practically zero (*K* ≫ *N*). Suppose we have selected two individuals following strategies *i* and *j*, so according to imitation dynamics, strategy *i* is replaced by strategy *j* in the population with probability

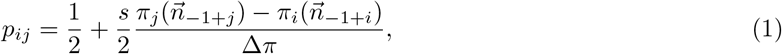

where 0 < *s* < 1 measures the strength of the selection, Δ*π* represents the highest payoff difference that can be realised in the game, while *π*_*i*_ and *π*_*j*_ denote the payoffs of *i* and *j* received previously by interacting with their completely different neighbours (Traulsen et al. (2005)). Since all players are randomly selected from the population, the expected change in the number of strategies *i* is calculated as

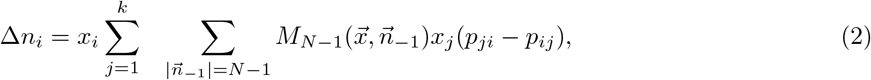

where 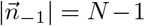 means that the summary is done for all cases where this equation is valid, 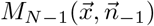 is the multinomial distribution with *x*_*i*_ global frequencies of the strategies 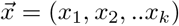. That is

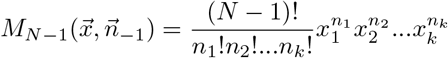

gives the probability that neighbours of a focal individual are present in the 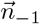 composition. Substituting *p*_*ij*_ and *p*_*ji*_ into (2), we get that

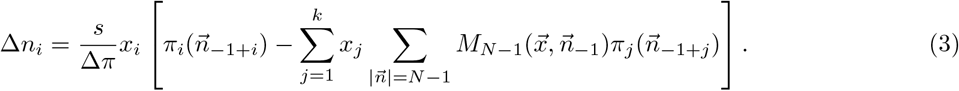

Assume that this elementary update is repeated many times, i.e. the number of updates is in the order of the population size, so that *x*_*i*_ the global frequency of strategy *i* changes according to the expected changes of strategy *i* in the population:

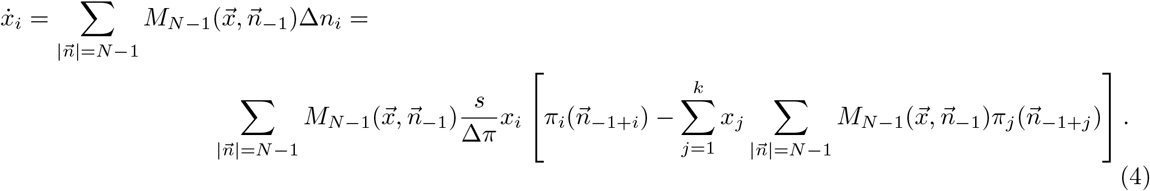

Neglecting the *s*/Δ*π* constant, which only rescales the time scale of the dynamics, and making trivial transformations, we obtain that

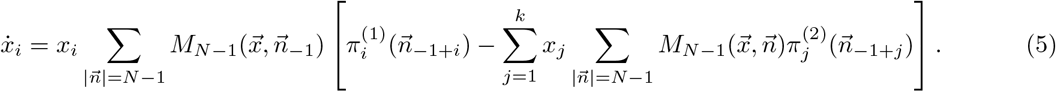

Introducing

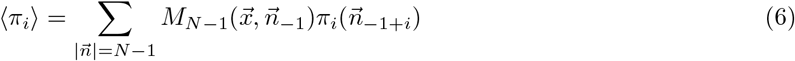

as the expected payoff of strategy *i* with *N* − 1 randomly selected members in the interaction group, and

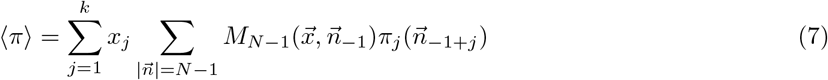

as the expected payoff in the population, we formally obtain the replicator dynamics

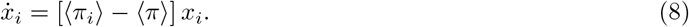

The important nature of this replicator dynamics is that the population level averages are computed in a slightly biased manner. Due to of the finite size of the local interaction groups the focal strategy is always overrepresented in the payoff averages (see e.g. 7). The correction term is of the order of 1/*N*, but as we will see later, this can have a crucial effect on the selection dynamics.

### 2.4. Application of the model: Public goods games

Let’s consider the example of an N-person public goods game where two strategies are defined, the cooperator strategy (*C*), which invests c units of energy in the public goods, and a defector strategy (*D*), which does not invest. The benefit (*b*) is determined by the total investment in the group, which is proportional to the number of cooperators (*n*_*C*_), thus

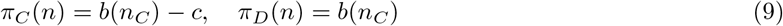

in a local neighbourhood, where *b*(*n*_*C*_) is an arbitrary function of *n*_*C*_. So strategy *D* always has a higher payoff than strategy *C* in a local neighbourhood. However, as we have shown, the *average* payoff of a cooperator and a defector are different even in a well-mixed population where interaction is local, i.e.

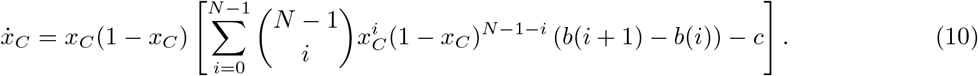

Depending on the functional form of *b*(*i*) and the maximal marginal benefit-cost ratio (*Max*[*b*(*i*+1)−*b*(*i*)]/*c*), many qualitatively different dynamics of (10) are possible (e.g. Hauert et al. (2006c); Archetti and Scheuring (2012)).

### 2.5. Linear Public Goods Game (PGG)

The classical public goods game assumes that the total investment of cooperators is summed and multiplied by the reward factor *r*, and that the total benefit is distributed equally to each individual. Thus, the payoffs of strategies *C* and *D* are

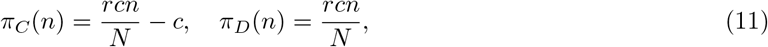

where *n* is the number of cooperators and *N* − *n* is the number of defectors in the group. Substituting the above payoffs into (10) we get

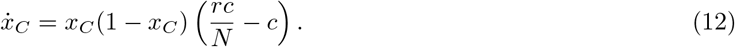

Thus cooperators win over defectors if *r*/*N* > 1, otherwise defectors win the selection (Hauert et al. (2006a)). Since in most biologically reasonable cases *r* < *N* (the reward is smaller than the number of individuals involved in the public good distribution), classical PGG leads to the fixation of defectors (Archetti and Scheuring (2012)).

### 2.6. Threshold Public Goods Game (TPGG)

The more general example is when the common good is a nonlinear s-shaped function of the number of cooperators in the interacting group. This biologically more relevant model (Archetti and Scheuring (2012)) is routinely approximated by the threshold dilemma game (Fig. 1) because the two models behave in qualitatively the same way, while the analysis of the TPGG is simpler (Archetti and Scheuring (2012, 2016)). According to the definition of the TPGG, if there are at least *k* number of cooperators among the *N* interacting individuals, then all of them receive the benefit *b* > 0 (without losing generality, we can assume that *b* = 1), but if the number of cooperators is below the threshold *k*, then there is no benefit, only the cooperators suffer the cost of cooperation (0 < *c* < 1), regardless of the actual number of cooperators among the interacting individuals (Table 1).

**Table 1:**
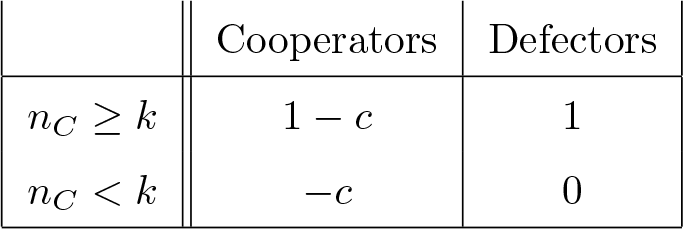
Payoffs of cooperators and defectors depending on whether there are enough cooperators in the interaction group (*n*_*C*_ *≥ k*) or not (*n*_*C*_ *< k*).

It follows from the structure of the payoff function (Table 1) that defectors always receive a higher payoff than cooperators. Thus, defectors win over cooperators in an infinite well-mixed population where there is no variance in the composition of strategies of interacting individuals, which practically means that the number of interacting individuals is very large. Our previously introduced model differs from this one solely in that the number of interacting individuals is not astronomical, so there is variance in their composition due to random selection.

Substituting the payoffs of the TPGG into (6), the average payoffs of *D* and *C* are

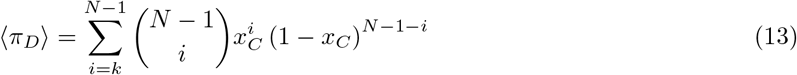

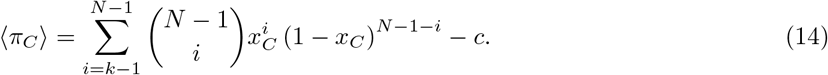

Consequently, the replicator dynamics of the system will be

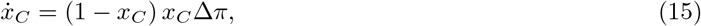

Where

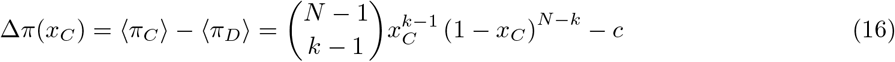

We note that this equation is identical to the replicator equation derived from the structured deme model (Archetti and Scheuring (2010)). There are two trivial fixed points of (15) the 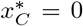 and 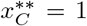. As we mentioned in the introduction, the most important property of TPGG in the structured deme model are that cooperators can coexist with defectors if the maximal marginal benefit of cooperation is greater than the cost of cooperation, i.e. if *Max*{Δ*π*(*x*_*C*_)} = Δ*π*_*Max*_ > 0 and 1 < *k* < *N*. We will present the complete analysis of the dynamical behavior of the system in the next section (but see Fig. 2).

## 3. Density dependent threshold public goods game

We have shown above that coexistence of cooperators and defectors in TPGG is possible in a population where there is no spatial aggregation or group formation, except that individuals that mix intensively in a large population interact with their local neighbours.

In this section we study the same model with the addition of a density dependent part of the population dynamics. Imagine that a population is present in a habitat with a maximal carrying capacity of *K*. In practice, this means that there are *K* places (or territories) available for individuals on the habitat. We compare two randomly selected individuals in the reproductive phase and the payoffs are calculated in the same way as before, but the offspring of the winner of the competition is randomly placed in a location of the habitat. If the selected location is empty, then the offspring has settled there; if it is occupied, then the replication was unsuccessful. In parallel, each individual dies with probability *d*. Replication and death cycles follow each other *n* times for a population of size *n*. These replication and death phases are followed by a payoff update phase as above for all individuals, and so on, replication, death and payoff collection cycles follow each other in the same order.

First we determine the dynamics of the whole population. Reproduction increases the size of the population if offspring is placed in an empty place, which happens with a probability of 1 −*n*/*K*. Furthermore, the rate of reproduction is proportional to the total density of the population *n*. (The replication rate doesn’t depend on the frequency of strategies, since there is one replication for each pairwise comparison). In parallel, each individual dies with probability *d* within a death cycle. So the dynamics describing the change of the total population in the deterministic limit is

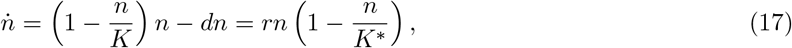

where *r* = 1 − *d, K*^*∗*^ = *K*(1 − *d*). The dynamics is identical to the well-known logistic equation, where the total population size converges to the stable fixed point *n*^*∗*^ = *K*^*∗*^ exponentially with speed proportional to *e*^*−rt*^. Consequently, after the transient phase with characteristic time 1/*r*, the population size stabilises at *n*^*∗*^, irrespective of the ratio of cooperators to defectors within the population.

Assume that the population already passed the transient phase, that is *n* ≈ *n*^*∗*^. Let *n*_*C*_ and *n*_*D*_ denote the number of cooperators and defectors in the population dynamical equilibrium (*n*_*C*_ + *n*_*D*_ = *n*^*∗*^). We have shown that the imitation dynamics used above leads to replicator dynamics in the deterministic limit, so we can consider equation (15) in the population dynamical equilibrium for the dynamics of strategies.

As we mentioned in the introduction the qualitative behaviour of (15) is well known (Archetti (2009); Archetti and Scheuring (2016), Fig. 2), which we summarize here point by point to compare the density independent model with the density dependent one:

- If *k* = 1 then there are three fixed points, the trivial 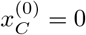 and 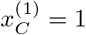, both of which are unstable, and the 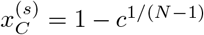 which is the globally stable fixed point of the system (Fig. 2A). Thus, if a single cooperator is sufficient to achieve high fitness, then the coexistence of cooperators and defectors is the only stable state of the dynamics.
- If 1 < *k* < *N* and Δ*π*_*Max*_ = [(*k* − 1)/(*N* − 1)]^*k*^[(*N* − *k*)/(*N* − 1)]^*N−k*^ − *c* > 0 then there are four fixed points, 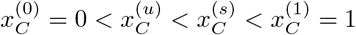.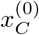 and 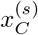 are stable fixed points with basin of attraction 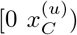 and 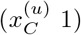 respectively, while 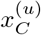 and 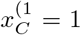 are unstable fixed points (Fig 2B). That is, if the cost of cooperation is below a critical level, cooperators and defectors will stably coexist if the initial frequency of cooperators is above 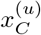, otherwise cooperators will be selected out.
- If *k* = *N* and Δ*π*_*Max*_ > 0 still true then there are three fixed points of the system 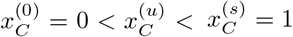. That is 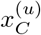 unstable fixed point separates two stable states where only defectors 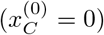 or only cooperators are present 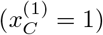 (Fig. 2C).
- If the above condition is not satisfied for the cost *c*, that is, if Δ*π*_*Max*_ ≤ 0 and *k* > 1 then the system has only the stable 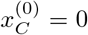 and the unstable 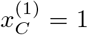 fixed points (Fig. 2D). This means that if the cost of cooperation is above a critical level, cooperators will always be selected out, regardless of their initial frequency.

Besides the parameters of the model, the dynamics of cooperators is only determined by the initial frequency of cooperators (see 15).

Notice that he role of empty sites, where there are no individuals, and defectors are identical in this sense; they don’t cooperate. Thus (15) remains valid in this density dependent version of the threshold dilemma game after the transient phase in population dynamics. However, due to the population dynamics *n*^*∗*^ = *K*(1 − *d*) in the dynamical equilibrium, so *x*_*C*_ = *n*_*C*_/*K* ≤ (1 − *d*). Thus, the equilibrium frequency of cooperators depends not only on the number of interacting individuals (*N*), the threshold (*k*) and the cost of cooperation (*c*), but also the decay rate (*d*) which determines a constraint on the maximal rate of cooperators in the habitat. Considering this constraint the possible dynamical behaviour of (15) is modified qualitatively:

- As mentioned above, there is only one stable fixed point of the replicator dynamics with 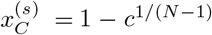 when *k* = 1. This 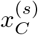 is the average probability of finding a cooperator in the *habitat*, so 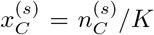. This means that the equilibrium frequency of cooperators is 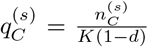 within the *population*, assuming that 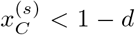. Consequently, as *d*, the mortality rate in the population increases, the frequency of cooperators in the population increases proportionally to 1/(1−*d*). However the frequency of cooperators continues to increase just until 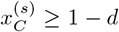. Above that *d* value 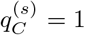, since then all individuals in the population should be cooperators in order to reach the possible maximum and evolutionarily stable fitness.
- Consider the case where 1 < *k* < *N* and Δ*π*_*Max*_ > 0. Assume that 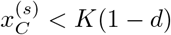 and 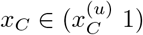 initially. Then, as we have shown above, the equilibrium ratio of cooperators in the habitat will be 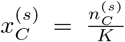. Thus, the frequency of cooperators in the population is 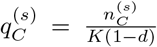 as in the previous case. Similarly, in the case when *d* ∈ [*d*_*cr*1_ *d*_*cr*2_], where *d*_*cr*1_ and *d*_*cr*2_ are determined by 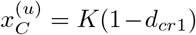 and 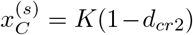, the replicator dynamics drive *x*_*C*_ to the maximum attainable frequency of cooperators, 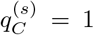. However, in the case when *d* > *d*_*cr*1_, the initial frequency of cooperators is always lower than 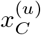, so *x*_*C*_ and consequently *q*_*C*_ converge to zero. Of course, the same happens if 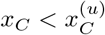 initially, regardless of the death rate *d*.
- If *k* = *N* and Δ*π*_*Max*_ > 0 is still valid, the system behaves similarly as in the previous case. Assume that 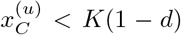 and 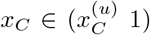 initially. Then, the equilibrium ratio of cooperators in the habitat will be 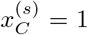. Thus, cooperators fixates in the population. However, if *d* > *d*_*cr*1_) then the initial frequency of cooperators is always lower than 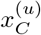, so *x*_*C*_ and consequently *q*_*C*_ converge to zero as before. Of course, the same happens if 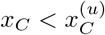 initially, regardless of the death rate d.
- If 1 < *k* ≤ *N* and Δ*π*_*Max*_ ≤ 0 then the cooperators will have always lower average fitness than defectors, regardless of population density and initial frequency of cooperators, so the only stable fixed point of the system is 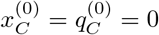.

## 4. Agent based models and simulations

### 4.1. The density independent models

One of our main aims of this section is to compare the dynamical behaviour of the model in the replicator dynamics limit with the results from the stochastic agent-based version of the model. We define agents being either cooperators or defectors in a population of *K* individuals (*K* = 2000 in most simulations). Agents play the TPGG as defined earlier with interacting group size *N* and threshold *k*. In the interaction phase two groups of individuals of size *N* are formed randomly. We then randomly select one individual from each of the two groups (let us denote them with *i* and *j*) and calculate their payoff *w*_*i*_ and *w*_*j*_, respectively. The probability of individual *i* being replaced by individual *j* is determined by (1), where *π*_*i*_ *p*_*ij*_ and Δ*w*_*max*_ are computed from the Table 1. This process is repeated *K*/2 times for a single Monte Carlo (MC) cycle, corresponding to one generation of updates.

Using this algorithm, we estimate the stable and unstable fixed points of the dynamics at different *N* and *k* values in the agent-based model and compare these values with the fixed points calculated from (15). The initial proportion of cooperators in the population was typically 0.5 (or even higher for higher *k*/*N* ratios), and their proportion is calculated at the end of each generation to track their frequency. The procedure continues until one of the strategies dies out or a polymorphic equilibrium state is reached after 200 generations. The mean frequencies of the strategies are calculated as the average of ten independent simulations where for each simulations the mean values of the last 100 generations were computed. To estimate the unstable fixed point we run a series of simulations with fixed *k* and *N* parameters and with different initial ratios of cooperators. We repeated the simulations ten times for each fixed parameter and initial value and counted the number of cases when the system stabilised in the polymorphic state and when it went into the pure defector state. The unstable fixed point is located at the initial value from which the dynamics is started, in half of the cases the system goes into the polymorphic state and in the other half of the cases into the defective state.

As the results of the simulations demonstrate, the estimated stable and unstable fixed points of the agent-based model, which is a numerical realisation of our well-mixed model system, closely approximate the fixed points of the deterministic dynamics in infinite population. The results confirm the correctness of the mathematical calculation presented above and also show that the infinite deterministic model can be well applied to relatively small populations (*K* = 2000) and even when the selection is not very strong (*s* = 0.6).

We also examined a variant of the agent-based model defined above, in which the number of interacting individuals fluctuates around an average due to a kind of stochasticity in the carrying capacity of the habitat. That is, the fluctuation is independent of the ratio of strategies, it is the consequence of density fluctuation present in every natural population. Fluctuation was incorporated into the model by randomly selecting the number of interacting individuals from a given interval. Thus, the actual size *N*_*l*_ of interacting groups is chosen uniformly from [*N* − *l, N* + *l*] interval (*N*_*l*_ ∈ {*N* − *l, N* − *l* + 1, .*N* + *l*} (*l* = 0, 1, 2,..); *N* − *l* > 1), so *V*_*l*_, the variance of *N*_*l*_, is 1/3*l*^2^.

We investigated how the equilibrium frequency of cooperators changes as the variance of the group size increases, while keeping the other parameters of the model constant. Figure 5 presents our findings, demonstrating that an increase in variance has a non-monotonic impact on the equilibrium cooperation level when *k* > 1. In fact, the equilibrium frequency of cooperators increases until reaching a certain level of variance, then decreases beyond this point. If one cooperator is sufficient for high fitness (*k* = 1), then increasing the variance of the interacting group size monotonically increases the equilibrium frequency of cooperators (Fig. 5 left upper subfigure).

**Figure 4.**
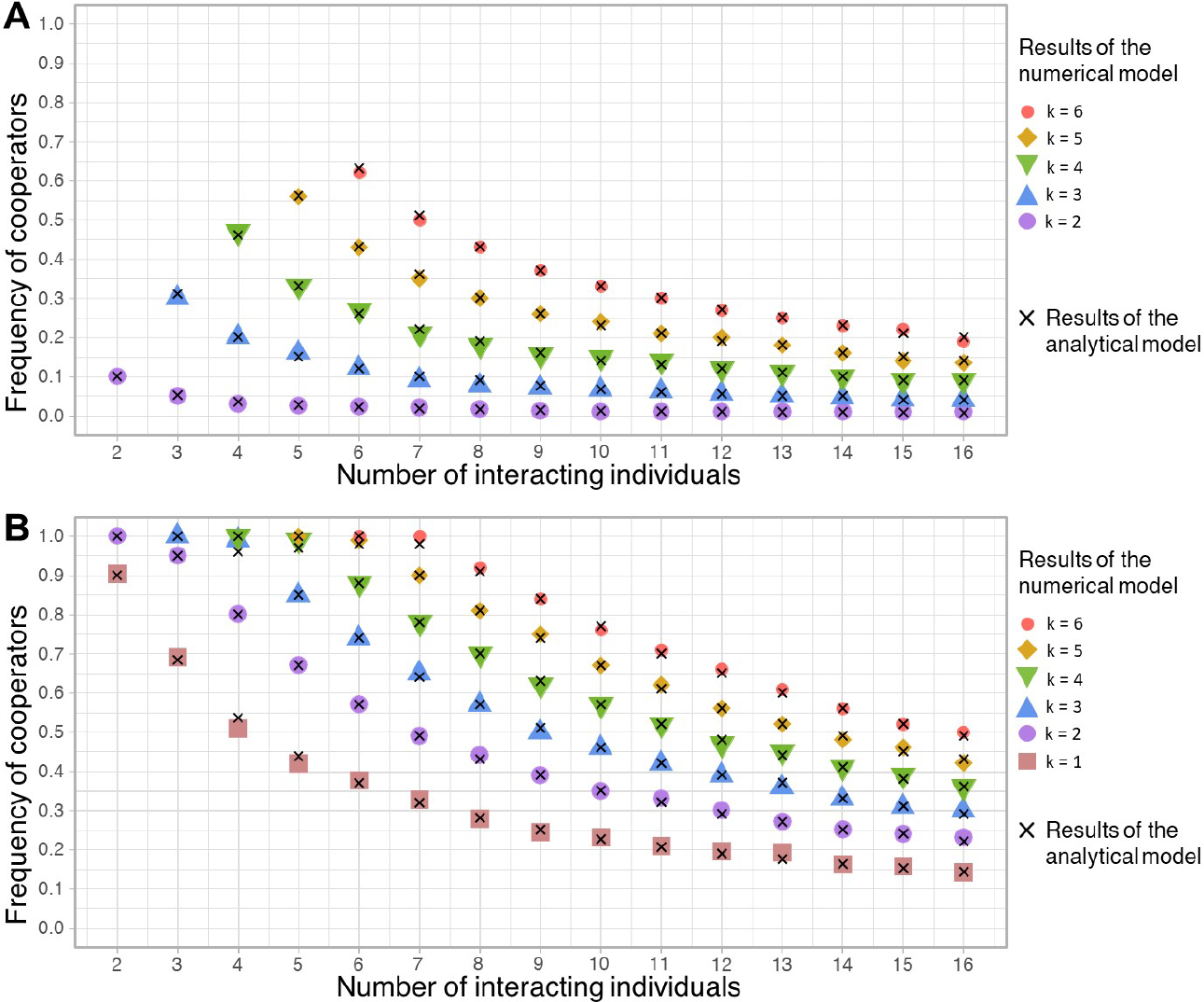
The unstable and stable fixed points in function of the interaction groups size and threshold value (*k*) according to the replicator dynamics and the agent-based model. **A:** The unstable fixed points. **B:** The stable fixed points. Crosses denote the fixed points computed from the replicator dynamics while the nearly identical measured fixed points of the agent based model are denoted by different filled symbols.

**Figure 5.**
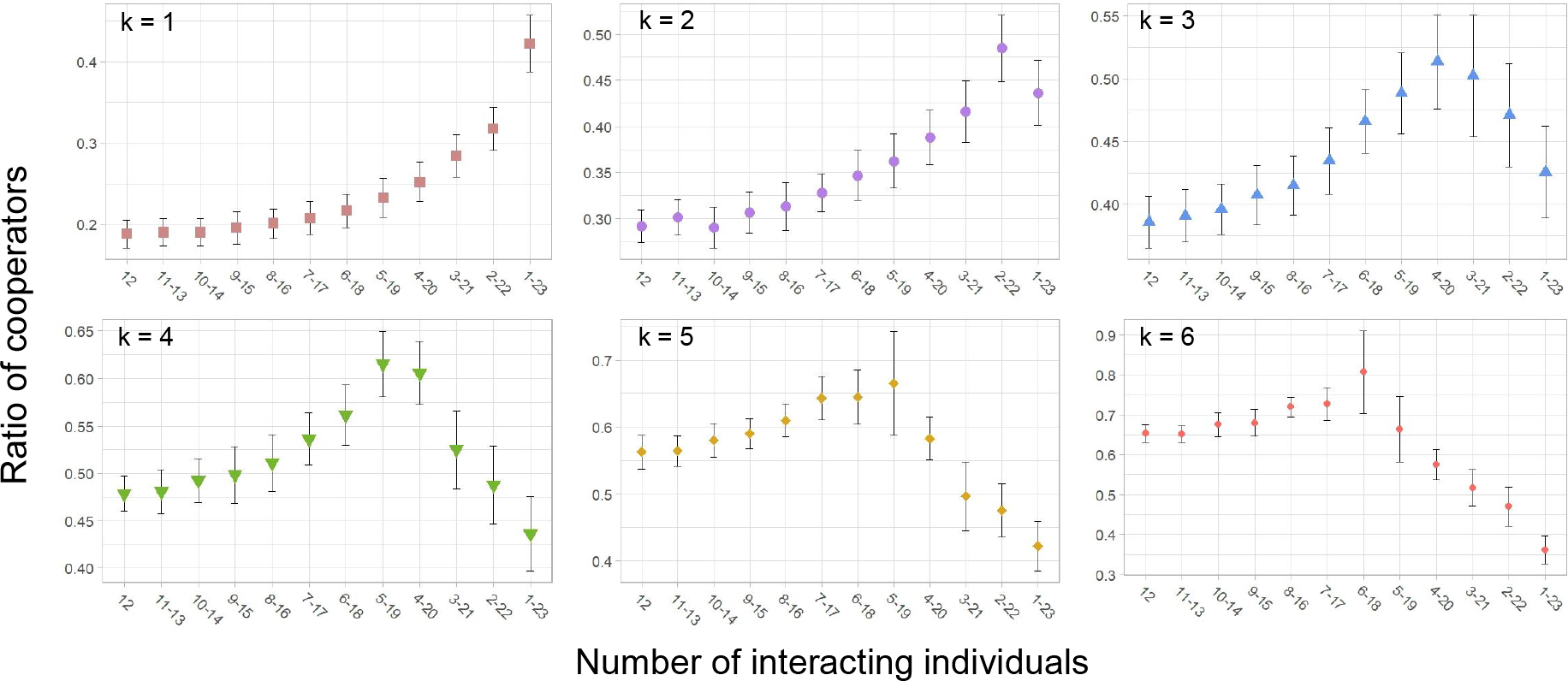
Effect of the variance in the size of the number of interacting individuals on the dynamics at different thresholds. The mean equilibrium frequency of cooperators and its variance are plotted as a function of the size of the interval from which the actual number of interacting individuals is randomly selected. Mean and variance are calculated from ten simulations. *K* = 2000, *s* = 0.6, *c* = 0.1, *N* = 12, *l* = 0, 1, 2,..11.

### 4.2. The density dependent model

To investigate the effect of density dependency we modified the density independent agent based model in three ways. First, in addition to updating strategies according to the rule (1), individuals die with probability *d* in each Monte Carlo cycle. Thus, although there are *K* discrete sites in the habitat, individuals do not occupy all of them. This means that the actual interacting group size can be less than *N* which implies the second differences: When interaction groups are formed, we randomly select *N* sites from the habitat, some of which may be empty sites due to mortality. Third, the winning strategy doesn’t replace the losing strategy in the population, but replaces its copy at a new randomly selected site. This replacement is only successful if the offspring is placed in a vacant site of the habitat. In the simulation, an MC cycle of competition and replication is followed by an MC cycle of death events, where an MC cycle is equal to the actual population size *n*. This algorithm is continued until one of the strategies disappears from the population or a polymorphic steady state is reached. Figure 6 depicts the time series of cooperator and defector strategies at characteristically different death rates which demonstrates that population reaches the equilibrium density within some dozens of generations which was our assumption in the analytical model. To reach the dynamical equilibrium of the strategies needs more time which can vary from hundred generations till thousand of them depending on the death rate.

**Figure 6.**
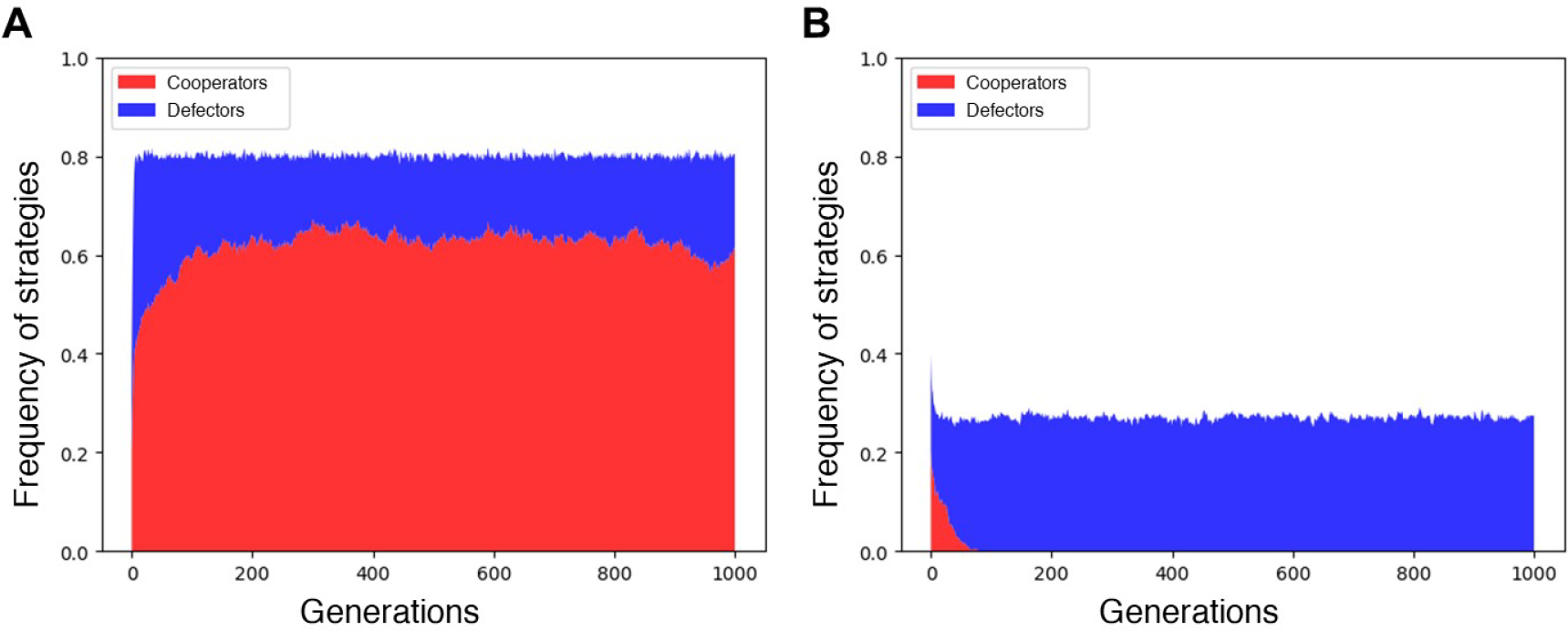
The population and frequency dynamics in the density dependent model. Population density reaches its equilibrium within some generations. After that short transient phase only the ratio of the strategies can change meaningfully. **A:** Stable coexistence of cooperators and defectors at lower death rate, (*d* = 0.2), **B:** Cooperators are selected out at high death rate, (*d* = 0.73). Other parameters are: *K* = 5000, *N* = 6, *k* = 3, *c* = 0.2, *s* = 0.6, initially *K/*2 sites are fulfilled by either cooperators or defectors with the same ratio.

Figure 7 depicts how the equilibrium frequency of cooperators changes as a function of population density measured by the death rate. The results of the simulations are in very good agreement with the results of the analytical model, demonstrating both the validity of the assumptions of the analytical model and the negligible effect of stochasticity and finite size in the numerical model.

**Figure 7.**
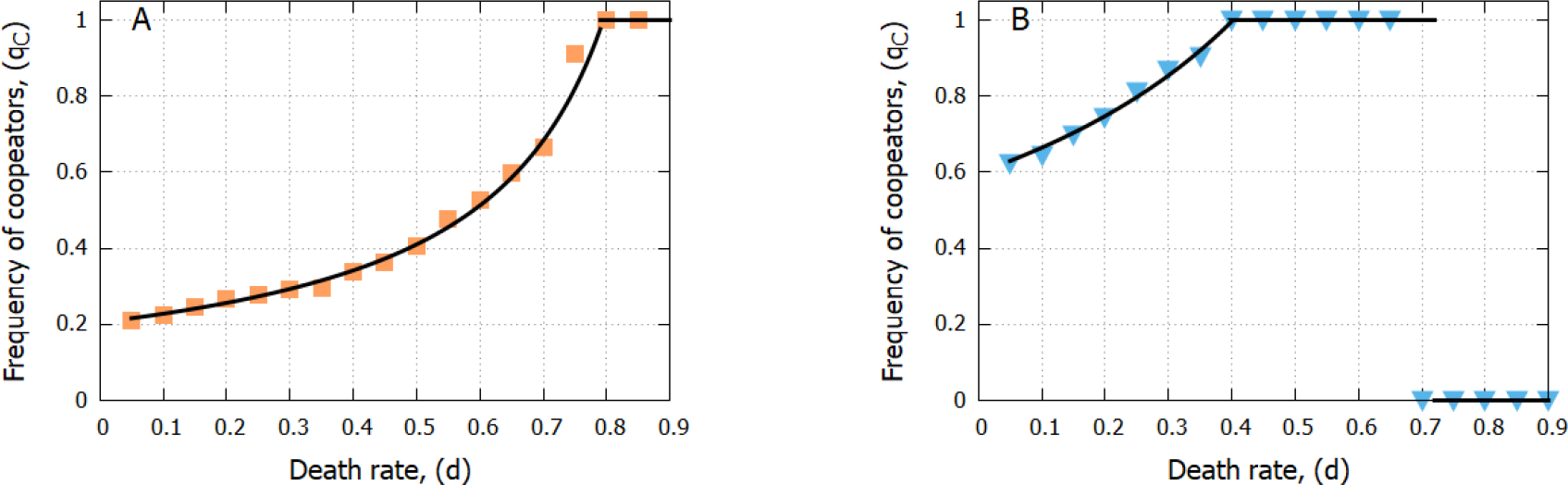
Equilibrium frequency of cooperators as a function of death rate. The solid line shows the predictions of the analytical model, the squares and triangles show the values obtained by simulating the agent-based stochastic model. **A:** *k* = 1 **B:** *k* = 4, other parameters are *K* = 5000, *N* = 8 *c* = 0.2, *s* = 0.6. Numerical simulations run for 1000 generations, frequencies are calculated as the average value of the last 100 generations.

## 5. Discussion

We have introduced a model of general N-person game where population is well mixed and there is no assortment. Individuals are compared in pairs and transmit their strategies to the next generation depending on their relative payoffs. We show that the dynamics of the strategies in the deterministic limit case are described by the same replicator dynamics as in the structured deme model, despite the absence of multi-level selection in our model. Consequently, cooperators and defectors are typically coexist in TPGG in this well-mixed population. This result is verified by the stochastic agent-based version of the analytical model (Fig. 4).

It is a crucial feature of the model that the compared individuals were previously in different interaction groups due to the large population size and intensive mixing. If the compared individuals are from the same or partially the same group, then the defectors are favoured by selection Hilbe (2011). That is, in this case, spatial aggregation does not help the cooperator, as it is in other cooperative dilemmas and population dynamics Nowak and May (1992); Számadó et al. (2008); Smaldino and Schank (2012); Czárán and Scheuring (2022). It is clear that game interactions and competition between individuals for resources do not occur in a completely uncorrelated manner in real populations due to viscosity, so that the real dynamics lie somewhere between Hilbe’s model Hilbe (2011) (game and competition within the same interaction group) and our present model (competition between individuals from different interaction groups). The development of such a more realistic model will be the aim of a forthcoming work.

We also investigated the behaviour of a density dependent version of the model. We show that the maximum carrying capacity of the habitat and the natural mortality rate jointly determine the equilibrium population density. We also show that the population converges to this equilibrium density regardless of the actual frequency of strategies, where classical density-independent replicator dynamics can be applied, with the difference that there may be fewer neighbours than the maximum in the interaction neighbourhood due to empty sites in the habitat. As a consequence, as density decreases (i.e. as mortality increases), the equilibrium frequency of cooperators will increase, and then below a well-defined density, cooperators will become fixed in the population. If more than one cooperator is needed to reach the threshold (*k* > 1), then lowering the density further will lead to a too sparse population where *k* individuals will almost never be near each other, so cooperators will suddenly die out. If *k* = 1, then only one cooperator is needed to reach the threshold, so at any low density, cooperators will always be the winners of the selection (Fig. 7), since they always get their high fitness (1 − *c*) regardless of the group composition, while this is hardly true for defectors. These results are interesting for two reasons: First, it can be tested in microbial systems under laboratory conditions whether a decrease in density does indeed cause an increase, fixation and then sudden disappearance of the abundance of cooperators. Second, the prediction that a population of a species in a poorer habitat will have a higher proportion of cooperators when playing NLPGG than in a richer habitat can be tested in the field. These implications also raise the possibility that an increase in the production of extracellular materials, as some common goods in microbial communities, could be a signal of a sudden disappearance of the production of these materials, so it can be an early warning signal of a regime shift in the functioning of the microbes. This signal differs from previously proposed and detected early warning signals in ecological systems, such as increasing variance, increasing autocorrelation or skewness or shift in variance spectra (Scheffer et al. (2009); Dakos et al. (2012)).

We show that not only population density, but also its variation, affects the equilibrium cooperator frequency in a non-trivial way. The effect of varying group size on the behaviour of N-person games has been studied in several previous publications (Peña (2012); Peña and Nöldeke (2016, 2018); Broom et al. (2019)). For example, it has been shown that if the difference in the payoff functions of cooperators and defectors is an increasing (decreasing) convex (concave) function of the number of cooperators, then an increase in the variance of the group size increases (decreases) the equilibrium cooperator frequency (Peña and Nöldeke (2016)). If these conditions are not met, then no such clear statements can be made about the effect of group size variance. The TPGG falls into this mathematically ambiguous category because the payoff difference is neither concave nor convex (Peña and Nöldeke (2016)). We have numerically investigated this mathematically complex but biologically relevant case. We show that when the possible minimum group size is not less than the threshold, increasing the variance increases the equilibrium cooperator frequency. This trend is reversed when the possible minimum group size is less than the threshold (Fig. 5). The intuitive explanation for this behaviour is as follows: Since increasing the interaction group size while holding the threshold constant decreases the frequency of equilibrium cooperators at a decreasing rate (Fig. 4), and since actual group sizes are chosen evenly around the average group size, increasing the fluctuation in group size will increase the frequency of cooperators more when the actual group size is smaller than the average group size than when the actual group size is larger than the average group size. Consequently, the frequency of cooperators will increase at selection equilibrium. As the variance increases, this trend continues until occasionally there can be so few individuals in the interaction group that the number of cooperative individuals can’t fail to reach the critical value *k*. This can only happen if the minimum possible group size is smaller than *k*. Then, as the variance continues to increase, the equilibrium cooperator frequency starts to decrease because there will be more and more groups where there cannot be enough cooperators to reach the threshold. Of course, this reversal of the trend does not hold for *k* = 1, since no additional cooperators are needed to achieve high fitness (Fig. 5 A).

It should be noted that the benefit is not distributed among participants in the TPGG model; thus, all participants receive the same benefit at all densities. However, this is not necessarily the case in real biological scenarios (e.g. food sharing), where lower density may result in higher benefit per individual if a sufficient number of cooperators are present. In the future, we aim to explore the behaviour of games of this type.

Although the relationship between the model and field observations is rather loose, it is interesting to note that cooperative breeding in birds is more common in more rugged Cornwallis et al. (2017) and more uncertain habitats Rubenstein and Lovette (2007); Jetz and Rubenstein (2011), which is entirely consistent with what we have seen in the model.

## Acknowledgement

I.S. was supported by the European Union’s Horizon 2020 research and innovation programme under grant agreement No 952914. English grammar and style are checked by DeepL Write beta 2.0.

## References

Archetti, M., 2009. Cooperation as a volunteer’s dilemma and the strategy of conflict in public goods games. J Evol Biol. 22, 2192–2200. doi:10.1111/j.1420-9101.2009.01835.x.

Archetti, M., Pienta, K., 2019. Cooperation among cancer cells: applying game theory to cancer. Nat Rev Cancer 19, 110–117. doi:10.1038/s41568-018-0083-7.

Archetti, M., Pienta, K.J., 1995. Complex cooperative strategies in group-territorial african lions. Science 269, 1260–1262. doi:10.1126/science.7652573.

Archetti, M., Scheuring, I., 2010. Coexistence of cooperation and defection in public goods games. Evolution, 1140–1148doi:10.1111/j.1558-5646.2010.01185.

Archetti, M., Scheuring, I., 2012. Game theory of public goods in one-shot social dilemmas without assortment. J Theor Biol 299, 9–20. doi:10.1016/j.jtbi.2011.06.018.

Archetti, M., Scheuring, I., 2016. Evolution of optimal hill coefficients in nonlinear public goods games. J Theor Biol. 406, 73–82. doi:10.1016/j.jtbi.2016.06.030.

Bednarz, J., 1988. Cooperative hunting harris’ hawks (parabuteo unicinctus). Science 239, 1525–1527. doi:10.1126/science.239.4847.1525.

Broom, M., Pattni, K., Rychtá?r, J., 2019. Generalized social dilemmas: The evolution of cooperation in populations with variable group size. Bull Math Biol 81, 4643–4674. doi:10.1007/s11538-018-00545-1.

Charlesworth, B., 1979. A note on the evolution of altruism in structured demes. The American Naturalist 113, 601–605. URL: http://www.jstor.org/stable/2460278.

Clutton-Brock, T.H., O’Riain, M.J., Brotherton, P.N., Gaynor, D., Kansky, R., Griffin, A.S., Manser, M., 1999. Selfish sentinels in cooperative mammals. Science 284, 1640–1644. doi:10.1126/science.284.5420.1640.

Cornwallis, C., Botero, C., Rubenstein, D., Downing, P., West, S., Griffin, A., 2017. Weak selection helps cheap but harms expensive cooperation in spatial threshold dilemmas. Nat Ecol Evol 1, 0057. doi:10.1038/s41559-016-0057.

Czárán, T., Scheuring, I., 2022. Weak selection helps cheap but harms expensive cooperation in spatial threshold dilemmas. Journal of Theoretical Biology 536, 110995. doi:10.1016/j.jtbi.2021.110995.

Dakos, V., Carpenter, S., Brock, W., Ellison, A.M., Guttal, V., Ives, A., Kéfi, S., Livina, V., Seekell, D.A., van Nes, E.H., Scheffer, M., 2012. Methods for detecting early warnings of critical transitions in time series illustrated using simulated ecological data. PLoS ONE 7, e41010. doi:10.1371/journal.pone.0041010.

Hamilton, W.D., 1964. The genetical evolution of social behaviour. i. J Theor Biol 7, 1–16. doi:10.1016/0022-5193(64)90038-4.

Hauert, C., Holmes, M., Doebeli, M., 2006a. Evolutionary games and population dynamics: maintenance of cooperation in public goods games. Proc Biol Sci. 273, 2565–70. doi:10.1098/rspb.2006.3600.

Hauert, C., Holmes, M., Doebeli, M., 2006b. Evolutionary games and population dynamics: maintenance of cooperation in public goods games. Erratum for: Proc Biol Sci. 2006, 273:2565-70. Proc Biol Sci 273, 3131–3132. doi:10.1098/rspb.2006.3717.

Hauert, C., Michor, F., Nowak, M.A., Doebeli, M., 2006c. Synergy and discounting of cooperation in social dilemmas. Journal of Theoretical Biology 239, 195–202. doi:10.1016/j.jtbi.2005.08.040. special Issue in Memory of John Maynard Smith.

Hilbe, C., 2011. Local replicator dynamics: A simple link between deterministic and stochastic models of evolutionary game theory. Bull Math Biol. 73, 2068–2087. doi:10.1007/s11538-010-9608-2.

Jetz, W., Rubenstein, D.R., 2011. Environmental uncertainty and the global biogeography of cooperative breeding in birds. Current Biology 21, 72–78. doi:10.1016/j.cub.2010.11.075.

MacNulty, D.R., Smith, D.W., Mech, L.D., Vucetich, J.A., Packer, C., 2011. Nonlinear effects of group size on the success of wolves hunting elk. Behavioral Ecology 23, 75–82. doi:10.1093/beheco/arr159.

Matessi, C., Jayakar, S., 1976. Conditions for the evolution of altruism under darwinian selection. Theoretical Population Biology 9, 360–387. doi:10.1016/0040-5809(76)90053-8.

Maynard Smith, J., Szathmáry, E., 1995. The major transitions in evolution. Freeman, Oxford.

Nowak, M., May, R., 1992. Evolutionary games and spatial chaos. Nature 359, 826–829. doi:10.1038/359826a0.

Okasha, S., 2006. Evolution and the Levels of Selection. Oxford University Press. doi:10.1093/acprof:oso/9780199267972.001.0001.

Patel, M., Raymond, B., Bonsall, M., West, S., 2019a. Crystal toxins and the volunteer’s dilemma in bacteria. J Evol. Biol 32, 310–319. doi:10.1111/jeb.13415.

Patel, M., Raymond, B., Bonsall, M.B., West, S.A., 2019b. Crystal toxins and the volunteer’s dilemma in bacteria. Journal of Evolutionary Biology 32, 310–319. doi:10.1111/jeb.13415.

Peña, J., 2012. Group-size diversity in public goods games. Evolution 66, 623–636. doi:10.1111/j.1558-5646.2011.01504.

Peña, J., Nöldeke, G., 2016. Variability in group size and the evolution of collective action. Journal of Theoretical Biology 389, 72–82. doi:10.1016/j.jtbi.2015.10.023.

Peña, J., Nöldeke, G., 2018. Group size effects in social evolution. Journal of Theoretical Biology 457, 211–220. doi:10.1016/j.jtbi.2018.08.004.

Rosenthal, A.Z., Qi, Y., Hormoz, S., Park, J., Hsin-Jung Li, S., Elowitz, M.B., 2018. Metabolic interactions between dynamic bacterial subpopulations. eLife 7, e33099. doi:10.7554/eLife.33099.

Rubenstein, D.R., Lovette, I.J., 2007. Temporal environmental variability drives the evolution of cooperative breeding in birds. Current Biology 17, 1414–1419. doi:10.1016/j.cub.2007.07.032.

Scheffer, M., Bascompte, J., Brock, W., Brovkin, V., Carpenter, S.R., Dakos, V., Held, H., van Nes, E.H., Rietkerk, M., Sugihara, G., 2009. Early-warning signals for critical transitions. Nature 461, 53–59. doi:10.1038/nature08227.

Smaldino, P., Schank, J., 2012. Movement patterns, social dynamics, and the evolution of cooperation. Theor Popul Biol. 82, 48–58. doi:10.1016/j.tpb.2012.03.004.

Számadó, S., Szalai, F., Scheuring, I., 2008. The effect of dispersal and neighbourhood in games of cooperation. J Theor Biol. 253, 221–227. doi:10.1016/j.jtbi.2008.02.037.

Traulsen, A., Claussen, J., Hauert, C., 2005. Coevolutionary dynamics: From finite to infinite populations. Phys. Rev. Lett. 95, 238701. doi:10.1103/PhysRevLett.95.238701.

West, S.A., Cooper, G.A., Ghoul, M.B., Griffin, A.S., 2021. Ten recent insights for our understanding of cooperation. Nat Ecol Evol 5, 419–430. doi:10.1038/s41559-020-01384-x.

West, S.A., Griffin, A.S., Gardner, A., 2007. Social semantics: altruism, cooperation, mutualism, strong reciprocity and group selection. Journal of Evolutionary Biology 20, 415–432. doi:10.1111/j.1420-9101.2006.01258.x.

Wilson, D.S., 1977. Structured demes and the evolution of group-advantageous traits. The American Naturalist 111, 157–185. URL: http://www.jstor.org/stable/2459987.

Yip, E., Powers, K., Avilés, L., 2008. Cooperative capture of large prey solves scaling challenge faced by spider societies. Proc Natl Acad Sci U. S. A. 105, 11818–11822. doi:10.1073/pnas.0710603105.

